# Similarity in Functional Connectome Architecture Predicts Teenage Grit

**DOI:** 10.1101/2023.02.23.529637

**Authors:** Sujin Park, Daeun Park, M. Justin Kim

**Affiliations:** Department of Psychology, Sungkyunkwan University, Seoul, South Korea; Center for Neuroscience Imaging Research, Institute for Basic Science, Suwon, South Korea

**Keywords:** grit, functional connectome, neurodevelopment, adolescence, fMRI

## Abstract

Grit is a personality trait that encapsulates the tendency to persevere and maintain consistent interest for long-term goals. While prior studies found that grit predicts positive behavioral outcomes, there is a paucity of work providing explanatory evidence from a neurodevelopmental perspective. Based on previous research suggesting the utility of the functional connectome as a developmental measure, we tested the idea that individual differences in grit might be, in part, rooted in brain development in adolescence and emerging adulthood (*N* = 64, 11-19 years of age). Our analysis showed that grit was associated with connectome stability across conditions and connectome similarity across individuals. Notably, inter-subject representational similarity analysis revealed that teenagers who were grittier shared similar functional connectome architecture with each other, more so than those with lower grit. Our findings suggest that gritty individuals are more likely to follow a specific neurodevelopmental trajectory, which may underpin subsequent beneficial behavioral outcomes.

**Statement of Relevance:** Maintaining consistent effort and passion for long-term, personally meaningful goals – often referred to as grit – is suggested to be associated with a wide range of positive outcomes such as academic achievement, career success and subjective well-being. Although grit has gained substantial amount of interest not only in the academia but also from the general population, only a handful of studies have examined its neural underpinnings. Here, we examined whether putative developmental measures using whole-brain functional connectivity patterns (i.e., functional connectome) explain individual differences in grit. Using publicly available developmental neuroimaging dataset ranging from early adolescence to emerging adulthood, we found that functional connectome stability within individuals and similarity between individuals uniquely explained self-reported grit. Confirmatory analyses demonstrated the existence of common neural representations shared among gritty teenagers, which were unveiled during movie-watching. These findings highlight that grit may be embedded in the functional connectome architecture during adolescence and emerging adulthood.

## Introduction

Grit, an intrapersonal character to persevere and have sustained passion for long-term goals amidst setbacks (Duckworth et al., 2007), has been suggested as one of the key predictors of an individual’s academic success and beyond. Grittier high school students tend to earn higher grade point average and are more likely to graduate; grittier teachers, nurses, salespeople are more likely to maintain their jobs (Duckworth et al., 2007; Eskreis-Winkler et al., 2014; Jeong et al., 2019; Robertson-Kraft & Duckworth, 2014). The beneficial effects of grit reach one’s psychological realm: a recent meta-analysis highlighted a positive association between grit and subjective well-being (ρ = .46) (Hou et al., 2022). Adolescence is especially relevant because grit is shown to be malleable during this period (Park et al., 2018), and youth grit may yield long-lasting positive outcomes (Eskreis-Winkler et al., 2014; Jiang et al., 2019; Tang et al., 2019). As such, a recent meta-analytic study on the relations between grit and achievement shows that nearly 38% of studies have focused on adolescence with the larger effect sizes than childhood and adulthood (Lam & Zhou, 2022).

To date, only a handful of studies investigated neural markers of grit and those using functional magnetic resonance imaging (fMRI) indicated the prefrontal cortex (PFC) as a key region (Wang et al., 2017) and its functional connections with the striatum (Myers et al., 2016). Supporting the functional involvement of the frontostriatal network, Wang et al (2018) showed that adolescents’ grit was associated with the gray matter volumes of dorsolateral PFC and putamen, providing neuroanatomical evidence for the role of grit in persistence and selfregulation (Moriguchi & Hiraki, 2013). In another study, grit in children was correlated with the shape of the nucleus accumbens (Nemmi et al., 2016), a major component of ventral striatum where its dopamine system has been robustly linked with effortful and instrumental choices for rewards (Salamone & Correa, 2012; Treadway et al., 2012).

However, these studies mostly relied on *a priori* regions-of-interest (ROIs) and were limited to resting-state conditions, which is an important caveat considering that movie-watching paradigms are known to exhibit better reliability (Meer et al., 2020). In addition, as the functional architecture of the brain undergoes significant reorganizations during adolescence and early adulthood (Power et al., 2010; Kelly et al., 2009), a network level analysis of moviewatching fMRI data would be well-suited for capturing the dynamic trajectory of brain development and individual differences in grit.

Meanwhile, several pieces of evidence implicate that grit might be related to more mature functional brain systems encompassing cognitive and affective networks. Although grit is conceptually distinct from cognitive abilities, pursuing long-term goals while dismissing momentary distractions do require engagement of higher-order functions such as self-regulation (Wolters & Houssain, 2015; Duckworth & Gross, 2014) for which PFC is known to be responsible (Casey et al., 2011; Casey & Caudle, 2013). Considering that the PFC – a region suggested to be representative of brain maturity (Dosenbach et al., 2010) – was highlighted in prior neuroimaging research on grit, it is possible that gritty individuals benefit from a more mature functional architecture. Furthermore, recent studies reported positive associations of grit and cognitive reappraisal (Kalia et al., 2022; Valdez & Datu, 2021), an adaptive emotion regulation strategy that involves PFC engagement across development (McRae et al., 2012). Taken together, being gritty might capitalize on balanced interaction and maturation of cognitive and affective control networks.

In recent years, functional connectome (FC) – a whole-brain map of functional connectivity patterns between pairs of different regions – has emerged as a useful neural feature in explaining and predicting behavioral variability (Finn et al., 2015; Horien et al., 2019). Kaufmann et al. (2017) demonstrated that FCs individualize markedly in adolescence allowing identification of individuals’ FC, and delay in such distinctiveness was observed in those with clinical symptoms. Another study noted that the uniqueness of FC profiles starts to exist as early as 12 years of age and had a negative relationship with mental disorder symptoms (Shan et al., 2022), placing adolescence in the foreground of brain developmental period.

Recently, a novel approach using FCs as features of brain development has been proposed (Vanderwal et al., 2021). According to this method, a more nuanced brain-behavior relationship could be captured by correlating 1) connectomes across conditions (e.g., task-based or resting-state functional runs) within individual (i.e., within-subject connectome stability) and 2) connectomes across individuals in each condition (i.e., between-subject connectome similarity). Put another way, within-subject connectome stability denotes how stable one’s FCs are across conditions, whereas between-subject connectome similarity is how similar one’s FC in a given condition is to those of others. An important assumption is that connectome stability and similarity reflect two maturational processes occurring hand in hand for optimized neural processing. That is, brain development might not only involve FCs being more stable across conditions within individual, but they might also represent normative or template-like functional connectivity patterns shared among individuals in each condition. Supporting this idea, greater connectome stability and similarity were associated with higher social skills in a sample of 6-21 year-olds (Vanderwal et al., 2021). In a similar vein, a recent study demonstrated that within-subject stability and between-subject similarity to high scorers in cognitive tasks predicted sustained attention and working memory abilities in adults, highlighting the potential utility of the features in behavioral prediction models (Corriveau et al., 2022)

Considering these methodological advances in using whole-brain functional connectivity patterns, this study tests the following hypotheses. First, we expected that gritty individuals’ FCs might be relatively consistent across runs (i.e., high stability) and similar with others in each condition (i.e., high similarity) – two proxies as brain developmental measures. Second, networks that contribute to the relationship between grit and connectome stability and similarity would be distributed across the brain, notably networks that support cognitive-affective functions. In addition, we sought to test the potential utility of FC generated from fMRI data that do not require grit-specific tasks (e.g., movie-watching) in identifying gritty individuals, which in turn would be able to shed further light onto the neural underpinnings of grit.

## Materials and Methods

### Participants

All data used in this study were drawn from the publicly available Healthy Brain Network (HBN) dataset (Alexander et al., 2017). The dataset houses multimodal brain imaging as well as behavioral phenotypic data from various communities across New York City. The recruitment of the study was largely based on the families with clinical concerns in their child and the study was approved by the Chesapeake Institutional Review Board. For participants younger than 18, written consent and written assent were acquired from the legal guardians and the participants. Written informed consent was obtained for those older than 18. The dataset can be downloaded at http://fcon_1000.projects.nitrc.org/indi/cmi_healthy_brain_network/.

Data from the HBN Releases 7 through 10 were used for the present study, as the Grit scale was added to the protocol from Release 7. Among participants whose chronological age was between 11 to 19, a total of 238 participants with grit scores were initially accessed. Participant data were collected in one of two sites located at the Citigroup Biomedical Imaging Center (CBIC) or the Rutgers University Brain Imaging Center (RUBIC). Individuals without T1-weighted images as well as all four functional runs were excluded (*n* = 23). We removed low quality data after manually inspecting the preprocessed output (*n* = 12) and excluded participants with excessive head motion (mean framewise displacement (FD) > 0.2 mm) to avoid confounding effects from movement in the scanner (*n* = 129). After excluding data with FC construction failures (*n* = 10), the remaining 64 participants were analyzed as our main sample (*N* = 64, 22 females; age range = 11.07-18.82 years (*M* = 14.78); *n* = 48 from CBIC, see *Supplemental Methods*, Fig. S1 for sample characteristics).

### Phenotypic Data

Phenotypic assessments used in this study were obtained on the first and third visit of the HBN schedule. As a main behavioral variable of interest, the 12-item Grit scale was used to measure perseverance of effort (e.g., “I have overcome setbacks to conquer an important challenge.”) and consistency of interest (e.g., “New ideas and projects sometimes distract me from previous ones.”) for long-term goals on a 5-point scale (Duckworth et al., 2007). Items for consistency of interest were reverse-coded and the average score for the two subscales were calculated. Total grit score was averaged after adding up all the items resulting in a range from one (not at all gritty) to five (extremely gritty).

To show that grit is related to positive behavioral variables and to conceptually distinguish from cognitive abilities such as IQ or attention, we utilized several related phenotypic assessments available in the dataset. First, the Screen for Child Anxiety Related Disorders (SCARED) (Birmaher et al., 1999) and Mood and Feelings Questionnaire (MFQ) (Messer et al., 1995) were used as self-reported behavioral measures for anxiety and depression. Self-reported version of the Columbia Impairment Scale (CIS) was used as a measure for global impairment across domains including school performance, interpersonal relationship, psychopathology and use of leisure time (Bird et al., 1993). Parent ratings on attention deficit and hyperactivity level assessed by the Strengths and Weaknesses of Attention-Deficit/Hyperactivity Disorder Symptoms and Normal Behavior Scale (SWAN) were used (Swanson et al., 2001).

Two additional behavioral phenotypes were used as control variables in our main analyses. Based on the previous research demonstrating a link between FC stability and similarity measures and social skills (Vanderwal et al., 2021), we sought to ensure that the observed grit-brain association from our analyses was not attributable to social skills. To this end, the Social Communication Questionnaire (SCQ), a parent measure with 40 yes-or-no items often used as a screening instrument for autism spectrum disorder (Chandler et al., 2007), was used. Higher score in this assessment indicates impaired social communication skills including a child’s body movements and language or gestures usage. Among those who completed the Grit scale, all but one had SCQ scores and were thus analyzed (*n* = 63). We also used a composite score of full-scale intelligence quotient (FSIQ) from Wechsler Intelligence Scale for Children, Fifth Edition (Wechsler, 2014) to ensure that the observed effect was not derived by general intellectual abilities. A subsample (*n* = 41) in the range of 6-17 years of age provided both grit and IQ scores.

### Image Acquisition

Magnetic resonance imaging (MRI) data were collected using a Siemens 3T Tim Trio at Rutgers University Brain Imaging Center (RUBIC) and a Siemens 3T Prisma at Citigroup Biomedical Imaging Center (CBIC) on the second visit schedule. Identical scan parameters were used for both centers: 3D anatomical T1-weighted MPRAGE image (224 slices, repetition time (TR) = 2500 ms, echo time (TE) = 3.15 ms, flip angle (FA) = 8°, slice thickness = 0.8 mm), functional runs with echo-planar imaging sequences (TR = 800 ms; TE = 30 ms; FA = 31°; slice thickness = 2.4 mm; field of view (FOV) = 204 mm; multi-band acceleration factor = 6; voxel size = 2.4 mm isotropic).

The four functional runs were administered in the following order: two consecutive resting-state runs of 5.1 minutes each (Rest1 and Rest2; 375 TRs each) and two movie-watching runs of 10 minute (a clip from “Despicable Me” (MovieDM); 750 TRs) and 3.47 minute (a short animated movie “The Present” (MovieTP); 250 TRs) length. In resting-state runs, participants were asked to keep their eyes open and focus on the fixation cross on the screen. The latter two naturalistic stimuli were played with sound. In the MovieDM condition, a clip of a DVD version of the movie *‘Despicable Me*’ was extracted where the main character unwillingly reads the three kids a bedtime story. A full-length of the short animation *‘The Present*’, story of a boy getting a puppy as a surprise gift from his parents, was shown in MovieTP. We present full scan length results as our primary main analysis. For completeness, results from a volume-matched analysis using truncated volumes in the three functional scans (Rest1, Rest2, MovieDM) to fit the shortest scan length of MovieTP (250 TRs) are reported separately.

### Image Preprocessing

All fMRI data were preprocessed using fMRIPrep 21.0.2 (Esteban et al., 2019). This includes blood-oxygen-level-dependent (BOLD) signal reference image estimation, head-motion estimation, slice-time correction, co-registration, resampling onto standard space (MNI PediatricAsym:cohort-1) and confounds estimation. To increase signal-to-noise ratio, the data were smoothed with a 4 mm Full-Width at Half-Maximum Gaussian kernel using *3dmerge* in AFNI (Cox, 1996; Cox & Hyde, 1997). The first four dummy volumes were discarded and confound regression was performed with 27 regressors (24 head motion parameters: three translational and three rotational deviations and their squares, six temporal derivatives and their squares; three mean tissue signals: global, cerebrospinal fluid and white matter). Finally, we set our high pass filter cut-off as 0.01 Hz and low pass filter cut-off as 0.10 Hz to remove the physiological noise in the data.

### Functional Connectome Construction

Fully preprocessed fMRI images were then transformed into FCs using *3dNetCorr* in AFNI (Taylor & Ziad, 2013). By providing the Shen atlas parcellation with 268 ROI masks (Shen et al., 2013) to the timeseries data of each functional run as input, we generated a 268 × 268 wholebrain functional connectivity matrix for each individual. The correlation coefficients from each pair of brain regions were Fisher *z*-transformed. As a result, four FCs (Rest1, Rest2, MovieDM, and MovieTP) were created for each participant.

### Within-subject Connectome Stability

The overall workflow of this study is presented in Figure 1. Every whole-brain FC matrix was left with an off-diagonal lower triangular part and was vectorized. This left us with 35,778 connections (i.e., edges) between two different brain regions (i.e., nodes). In order to calculate FC stability within individuals, functional connectivity value of each edge was correlated across two conditions. Such edge-wise correlation resulted in three within-subject connectome stability measures: cross-rest (Rest1 and Rest2), cross-state (MovieDM and Rest1), and cross-movie (MovieDM and MovieTP) stability (Fig. 1*A*). Mean stability for each individual was calculated by averaging the three stability measures.

**Fig. 1.**
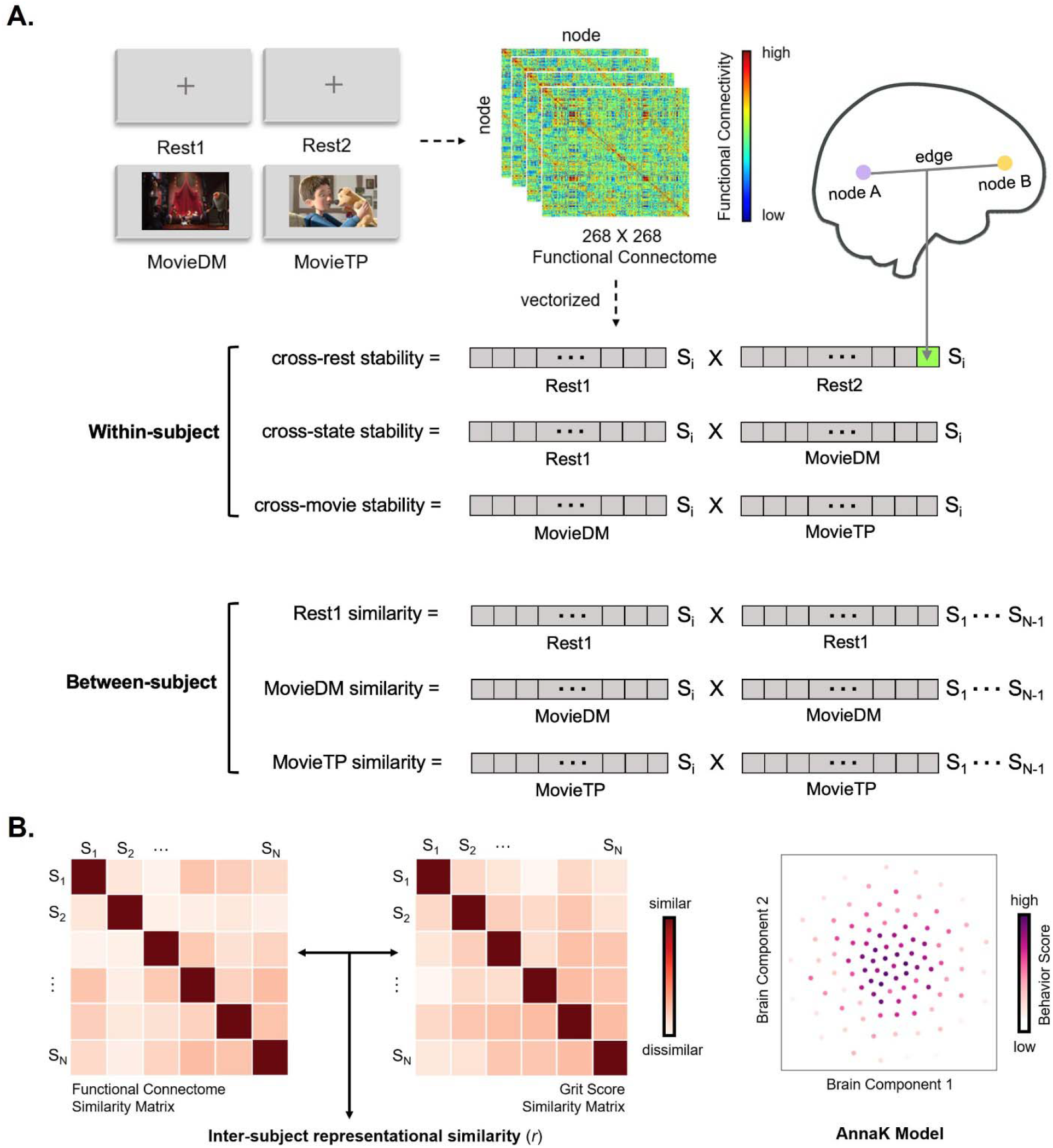
Summary of the analysis steps. **A.** Stability and similarity measures were created after vectorizing FCs of four conditions. Within-subject stability was calculated by correlating every functional connection in two different conditions for each subject (e.g., S_i_). Between-subject similarity in a given condition was the average of all edgewise functional connectivity correlations of one subject and all others. This procedure resulted in a total of six FC features (three stability and three similarity measures) for each subject. **B.** IS-RSA adopting the AnnaK model was conducted as a confirmatory analysis by comparing similarity matrices of brain feature (i.e., between-subject FC) and behavior variable (i.e., grit). According to the AnnaK model, high scorers in grit would share similar brain features more so than those with lower scorers.

### Between-subject Connectome Similarity

FC similarity across individuals in a given condition (i.e., MovieDM, MovieTP, Rest1) was computed by averaging edge-wise correlation coefficients between one participant’s FC and everyone else’s. In other words, FC in each run was correlated with FCs of all others excluding oneself. This resulted in three between-subject connectome similarity measures: Rest1, MovieDM, and MovieTP FC similarity (Fig. 1*A*). The three similarity measures were averaged to calculate mean similarity.

### Linking Functional Connectome Stability and Similarity with Behavior

Linear regression analysis was conducted to test whether the six FC measures (three stability measures and three similarity measures) predicted behavioral variables of interest (i.e., grit, social skills, IQ). Participants’ age, sex, scan site, and head motion for the corresponding functional runs (e.g., average mean FD of MovieDM and MovieTP for cross-movie stability) were entered as covariates. To investigate FC stability and similarity at the network level, we correlated grit and FC measures computed from predefined 10 canonical networks (Finn et al., 2015; Noble et al., 2017) separately.

### Inter-subject Representational Similarity Analysis (IS-RSA)

In addition to the main analyses using FC stability and similarity, IS-RSA (Finn et al., 2020) was conducted to confirm the relationship between FC similarity and grit. Specifically, via IS-RSA, we adopted 1) a nonparametric Mantel test where permutation-based null distributions of correlation values are used to determine significance (Mantel, 1967), and 2) a computational lesion approach whereby the relative contributions of canonical networks are quantified by creating a “virtual lesion” that knocks out specified edges per iteration. As IS-RSA allows flexible modeling of brain-behavior similarities, we assumed that gritty individuals may share more similar FCs (i.e., express normative functional connectivity patterns while in the same fMRI condition) with other gritty individuals, while those who are not may exhibit heterogenous FCs that are less similar to everyone else.

The idea of the IS-RSA framework is to compare the brain similarity matrix and behavioral similarity matrix (Fig. 1*B*). To this end, pairwise Pearson correlations of FC between one and every other subject were used to create a 64 × 64 inter-subject brain similarity matrix. For grit similarity, individuals were ranked by grit score from low (e.g., Rank 1 and 2) to high (e.g., Rank 63 and 64). The mean of the two individuals’ rank was regarded as a behavioral similarity metric for the pair (e.g., behavioral similarity for subjects Rank 63 and 64 is 63.5, Rank 1 and 2 is 1.5) resulting in a 64 × 64 inter-subject behavioral similarity matrix. This approach, called ‘the Anna Karenina (AnnaK)’ model, assumes that high scorers on a given behavior show higher brain feature similarity with each other whereas low scorers exhibit more variability (Finn et al., 2020) (Fig. 1*B*). Then, we computed inter-subject representational similarity using Spearman correlation between brain and behavior similarity matrices. The corresponding *p*-value was derived from the Mantel test with 5,000 permutation test iterations. The same procedure was conducted for other behavioral phenotypes that were correlated with grit as control analyses.

To check if there was a particular functional network contributing to the correlation between FC similarity and grit, the 10 canonical networks (Finn et al., 2015; Noble et al., 2017) were computationally lesioned. In this approach, in an iteration of the analysis, all nodes from a given network were excluded from the whole-brain FC to simulate a virtual lesion. If a network made a unique contribution in the prediction, we would expect a marked decrease in the intersubject representational similarity recalculated without the said network.

## Results

### Correlations Between Grit and Other Behavioral Measures

Correlations between grit and developmentally relevant behavioral measures are presented in Table 1. Grit had a significant negative relationship with anxiety (*r* = − .30, *p* = .02) and marginally significant relationship with depression (*r* = − .24, *p* = .06). Grit was also inversely correlated with general impairment (*r* = − .42, *p* < .001) indicating gritty teenagers tend to be less anxious and less susceptible to global impairment. In terms of social skills, there was a significant negative association (*r* = − .27, *p* = .03, such that gritty teenagers tended to possess higher social communication skills. Neither attention deficit scores nor IQ were correlated with grit (*r* = − .20, *p* = .12; *r* = .02, *p* = .92, respectively).

**Table 1.** Descriptive Statistics and Correlations for Study Variables.

| Variable | *M (SD)* | 1 | 2 | 3 | 4 | 5 | 6 | 7 |
| --- | --- | --- | --- | --- | --- | --- | --- | --- |
| 1. Grit | 3.07 (0.53) | - |  |  |  |  |  |  |
| 2. SCARED | 22.92 (15.29) | -.30* | - |  |  |  |  |  |
| 3. MFQ (*n* = 61) | 13.23 (12.61) | -.24 | .75** | - |  |  |  |  |
| 4. CIS | 11.16 (7.90) | -.42** | .49** | .57** | - |  |  |  |
| 5. SWAN | 0.32 (0.87) | -.20 | -.04 | -.06 | .24 | - |  |  |
| 6. SCQ (*n* = 63) | 8.05 (4.28) | -.27* | .04 | .15 | .23 | .44** | - |  |
| 7. WISC FSIQ (*n* = 41) | 104.71 (17.85) | .02 | .16 | -.09 | .11 | -.24 | -.36* | - |

*Note.* *N* = 64. In the case of a subsample, the number of participants is noted. *Abbreviations:* SCARED, Screen for Child Anxiety Related Disorders; MFQ, Mood and Feelings Questionnaire; CIS, The Columbia Impairment Scale; SWAN, The Strength and Weakness Assessment of Normal Behavior Rating Scale for ADHD; SCQ, Social Communication Questionnaire; WISC FSIQ, Wechsler Intelligence Scale Full Scale IQ.
\* $p < .05$ , \*\* $p < .01$ .

### Functional Connectome Stability and Similarity

As a summary measure of FC stability and similarity, we present the results based on mean stability and mean similarity. Group-average of mean stability was higher than mean similarity (full volume, mean stability = .45, mean similarity = .34; volume-matched, mean stability = .25, mean similarity = .19). Importantly, mean stability and mean similarity showed high positive correlation even after controlling age, sex, head motion and scan site (*r* = .64, *p* < .001) and this relationship remained when volume-matched (*r* = .49, *p* < .001). That is, individuals who showed highly stable functional connectivity patterns across conditions were more likely to resemble others’ FCs, replicating previous work by Vanderwal and colleagues (2021) and supporting the proposition that connectome stability and similarity may serve as concurrent brain development proxies. Stability and similarity measures were not correlated with age (*Supplemental Results*, Table S1), also replicating previous findings (Vanderwal et al., 2021).

### Functional Connectome Features and Grit

Partial correlation values from linear regression are reported in Table 2. Cross-movie stability and MovieTP similarity significantly predicted grit after controlling age, sex, scan site, and head motion (cross-movie stability, *r* = .34,*p* < .008; MovieTP similarity, *r* = .37,*p* = .003; Bonferroni corrected, *p* < .008). We also note that the significant positive relationship was largely driven by one of the two subscales of grit: perseverance of effort (cross-movie stability, perseverance of effort, *r* = .30, *p* = .02, consistency of interest, *r* = .23, *p* = .07; MovieTP similarity, perseverance of effort, *r* = .36, *p* = .005, consistency of interest, *r* = .23, *p* = .08). When the volumes were truncated to match the scan duration of the MovieTP condition, cross-movie stability was no longer a significant predictor (*r* = .16, *p* = .21). Therefore, as the FC similarity in MovieTP was the only robust predictor of grit when holding the scan length constant, we chose to focus on MovieTP similarity in subsequent analyses.

**Table 2.** Functional Connectome Features and Their Correlations with Grit Using Full and Truncated TRs.

| Variable | Within-subject FC Stability |  |  | Between-subject FC Similarity |  |  |
| --- | --- | --- | --- | --- | --- | --- |
|  | cross-rest | cross-state | cross-movie | Rest1 | MovieDM | MovieTP |
| Full volume |  |  |  |  |  |  |
| Grit | .03 (.8266) | - .09 (.5036) | **.34 (.0076)** | .04 (.7353) | .18 (.1785) | **.37 (.0032)** |
| Perseverance of effort | .01 (.9569) | .06 (.6725) | .30 (.0211) | - .04 (.7825) | .03 (.8195) | **.36 (.0045)** |
| Consistency of interest | .03 (.7933) | - .17 (.1949) | .23 (.0729) | .09 (.4776) | .22 (.0895) | .23 (.0812) |
| Volume-matched |  |  |  |  |  |  |
| Grit | - .09 (.5133) | .04 (.7823) | .16 (.2115) | .19 (.1498) | .21 (.1133) | - |
| Perseverance of effort | - .11 (.4219) | .16 (.2321) | .25 (.0545) | .08 (.5257) | .13 (.3108) | - |
| Consistency of interest | - .03 (.8011) | - .08 (.5502) | .02 (.8513) | .20 (.1345) | .18 (.1671) | - |

*Note.* Partial correlation coefficients ( $r$ ) between grit (total score and its subscales) and stability/similarity measures, controlling for age, sex, scan site and head motion are reported. Corresponding $p$ -values are in parenthesis and significant correlations in bold (Bonferroni corrected, $p < .008$ ).

In the MovieTP condition where FC similarity successfully predicted grit, none of the 10 networks was significantly correlated with total grit score by themselves (*Supplemental Results*, Table S2) implicating that simultaneously considering all of the networks offered better predictive performance than any single canonical network.

### IS-RSA

Based on the findings showing that grittier individuals shared more similar FCs with others in the MovieTP condition, IS-RSA was conducted as a confirmatory analysis to formally examine the shared geometries between FC and grit (Fig. 2). Correlation between MovieTP FC similarity and grit similarity matrices was significant (*r* = .28, *p* = .002) (Fig. 2*A*), confirming the FC similarity-grit association from the previous analysis. We then ran the same analyses using the subscales of grit and found that perseverance of effort and consistency of interest both showed similar results (perseverance of effort, *r* = .22, *p* = .02; consistency of interest, *r* = .21, *p* = .03). As the present IS-RSA adopted the AnnaK model, this implies that gritty individuals shared normative functional connectivity patterns with other gritty individuals while those who are not exhibited heterogenous FCs that are less similar to everyone else (Fig. 2*B*).

**Fig. 2.**
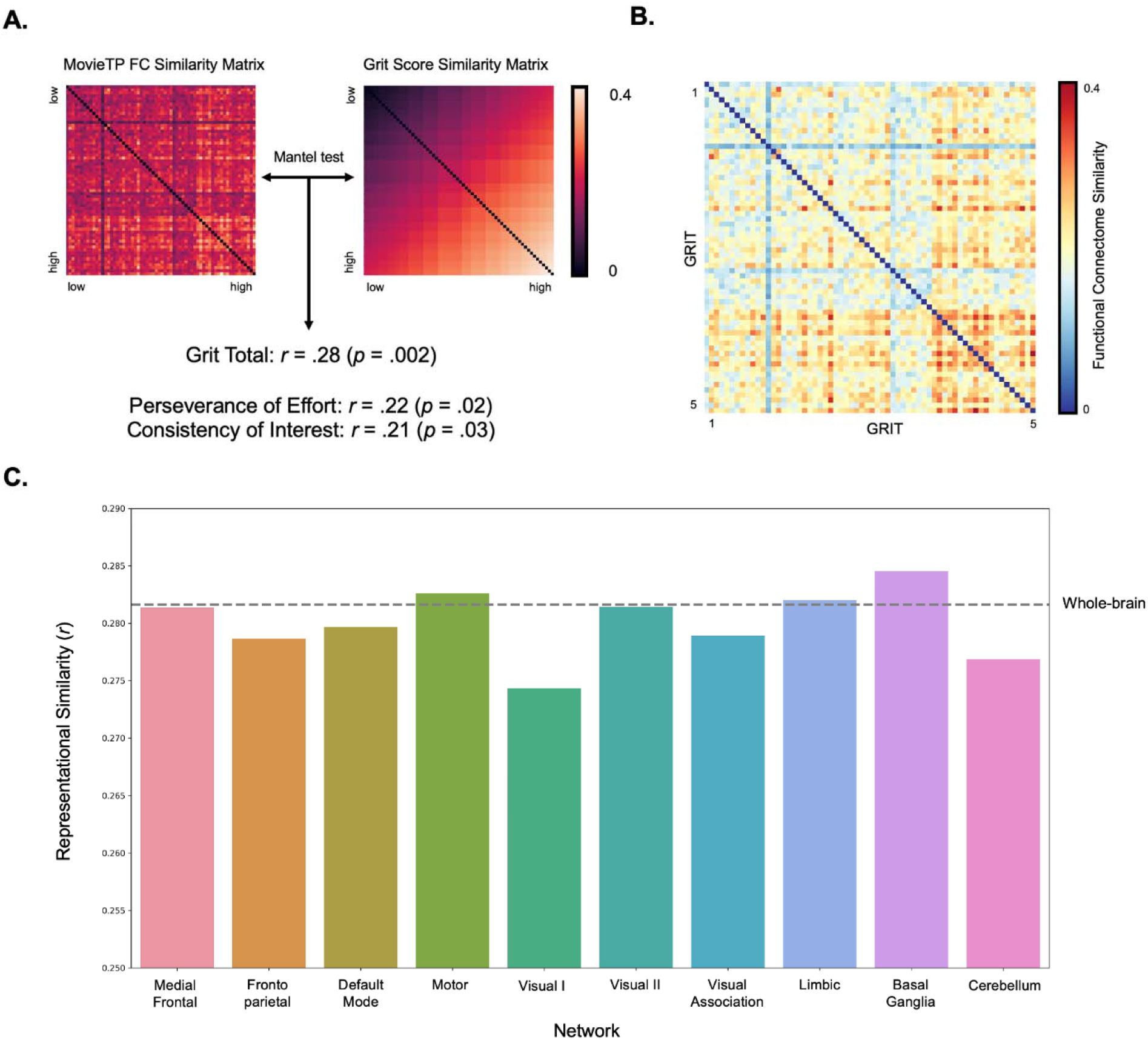
Results of the IS-RSA. **A.** MovieTP FC similarity (left) and grit score similarity (right) matrices were compared (5,000 permutation test iterations). Composite score of grit yielded higher *r*-value than its subcomponen**ts. B.** Visual summary of FC similarity sorted by grit score (from low to high). Gritty individuals on the bottom right section showed higher FC similarity among themselves. **C.** Results of the computational network lesion approach. Larger drop in inter-subject representational similarity (*r*) of each network lesion compared to that of the wholebrain result (dashed line) is interpreted as higher contribution of a network in IS-RSA. Single network lesions did not show significant drop-off in representational similarity (Bonferroni corrected, *p* < .005).

Results of the computational lesion approach suggest that, once again, the entirety of whole-brain networks, rather than any given single network, contribute more to the observed grit-brain relationship. There was no significant decline in correlation for any of the network lesions (Medial frontal, *r* = .28, *p* = .003; Fronto-parietal, *r* = .28, *p* = .002; Default mode, *r* = .28, *p* = .003; Motor, *r* = .28, *p* = .002; Visual I, *r* = .27, *p* = .003; Visual II, *r* = .28, *p* = .003; Visual association, *r* = .28, *p* = .002; Limbic, *r* = .28, *p* = .004; Basal Ganglia, *r* = .28, *p* = .003; Cerebellum, *r* = .28, *p* = .004; Bonferroni corrected, *p* < .005) (Fig. 2*C*). This finding may imply that predictive brain systems of grit do not rely on a single network, but rather they are distributed across the brain.

### Control Analyses

We performed a set of control analyses to validate the main findings that grit is related to neurodevelopmental features (Fig. 3). First, considering that most of the participants received clinical diagnoses at the point of data collection (Fig. 3*A*), it is possible that gritty individuals may have been different from less gritty individuals in terms of the prevalence of clinical conditions. Thus, a chi-square test was carried out to test whether there was a significant association between grit and clinical diagnoses. We split participants in half according to the median grit score (3.00) resulting in a high grit group (*n* = 31) and low grit group (*n* = 33). The number of participants with diagnoses (e.g., attention deficit hyperactivity disorder, anxiety disorders, major depressive disorders, cognitive abilities-related disorders, autism spectrum disorders) were not significantly related to grit scores (*X^2^* (6, *N* = 64) = 7.98, *p* = .24). Additionally, we provide group-averaged brain maps of functional connectivity patterns for high and low grit, which suggests that both groups had similar nodes exhibiting the strongest connections in the MovieTP condition (*Supplemental Results*, Fig. S2).

**Fig. 3.**
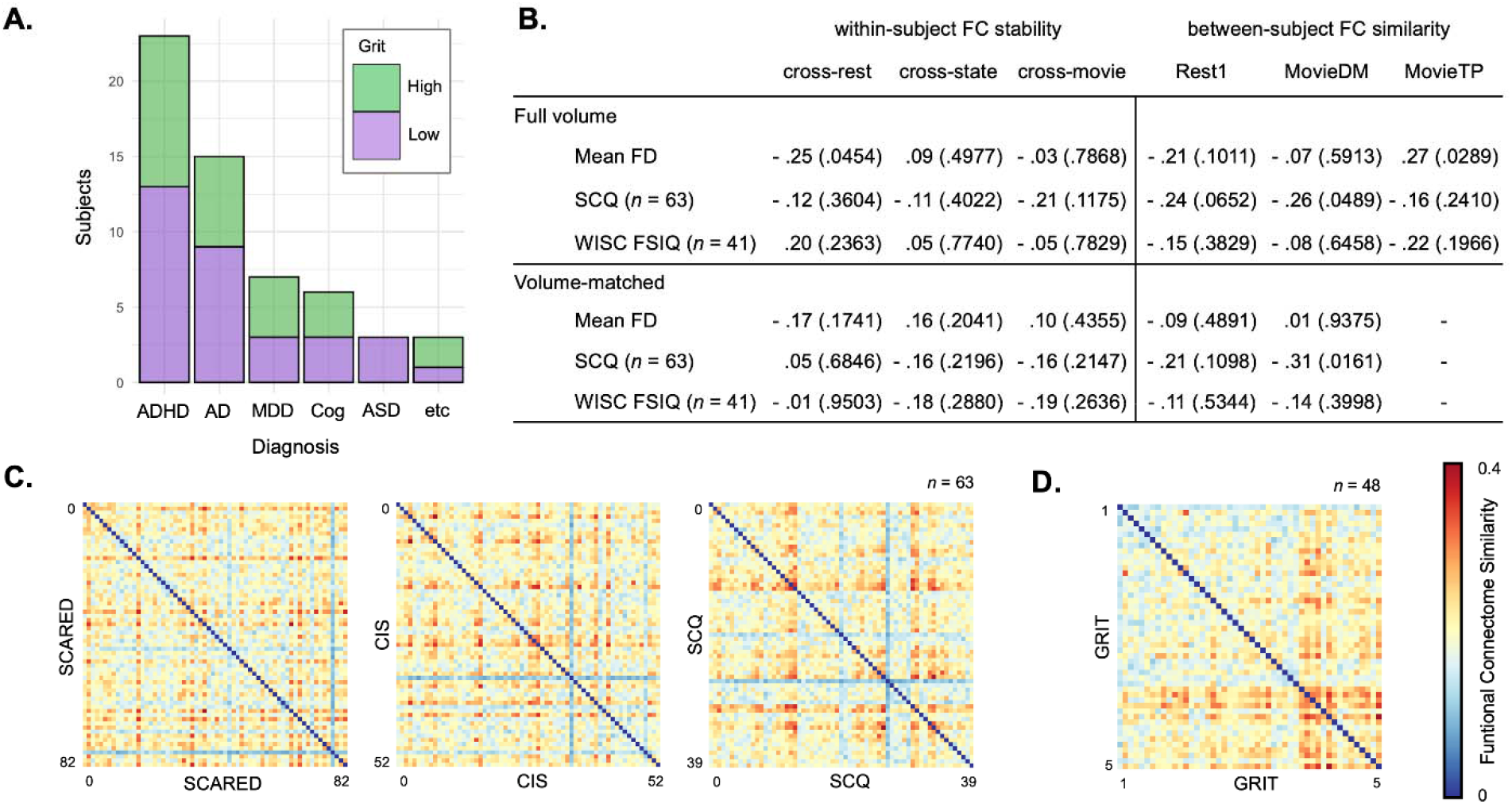
Summary of control analyses. **A.** Stacked bar graph of the participants’ clinical conditions, grouped by grit scores (median-split). *Abbreviations*: ADHD, Attention-Deficit Hyperactivity Disorder; AD, Anxiety Disorder; MDD, Major Depressive Disorder; Cog, Cognitive-related disorder; ASD, Autism Spectrum Disorder. **B.** Partial correlation coefficients (*r*) between control variables (Mean FD, SCQ, WISC FSIQ) and FC features in full length and volume matched analyses (Bonferroni corrected, *p* < .008). **C.** Heatmaps where FC similarities are sorted by behavior scores indexing anxiety, global impairment and social skills, respectively. Unlike grit, all three heatmaps do not show any discernable pattern. **D.** Heatmap depicting that gritty individuals share FC similarity with others, based on IS-RSA results from a subsample of CBIC participants.

Next, due to potential confounding effects of head motion during scans, we examined the correlation between head motion and the six measures of FC stability and similarity, where head motion was calculated by averaging the mean FD in the corresponding conditions. FC measures were not significantly correlated with head motion neither in the full volume nor in the volume-matched analyses (Fig. 3*B*), which may be attributed to conservative head motion threshold applied in this study.

To test whether the observed effects of the main results were specific to grit, the six FC measures were correlated with social skills and IQ while controlling for the effects of age, sex, scan site and head motion. Social communication skills had overall negative correlations (i.e., higher score in SCQ is regarded as lower social communication skills), but none of them survived the correction for multiple comparisons. Results of 41 participants with whom had both grit and IQ scores also showed that the six brain features were not predictive of IQ (Fig. 3*B*). In addition, when IS-RSA was conducted using other phenotypic data that were correlated with grit, including anxiety, global impairment, and social communication skills, there were no significant associations found for any of these measures with MovieTP similarity (SCARED, *r* = − .04, *p* = .65; CIS, *r* = − .13,*p* = .17; SCQ, *r* = − .08,*p* = .40) (Fig. 3*C*). These results provide supporting evidence for a unique association between MovieTP similarity and grit when tested with the AnnaK model.

Finally, even though the neuroimaging data collected in the HBN dataset followed identical scan parameters, MRI scanners from the two sites might yield different results. To address this possibility, an additional IS-RSA was performed in which the data were restricted to the 48 participants from CBIC. Results still showed that gritty individuals do share similar FCs with one another (*r* = .33,*p* = .001) (Fig. 3*D*), consistent with the results from IS-RSA with the full sample (Fig. 2*B*).

## Discussion

The purpose of the current study was to investigate whether brain development measures derived from FCs capture individual differences in teenage grit. Here, we showed that grit was significantly associated with movie-related within-subject FC stability and between-subject FC similarity. When matched for scan duration, FC similarity in the MovieTP condition was the sole predictor of grit, meaning that gritty teenagers showed more similar whole-brain FCs with others. IS-RSA with the AnnaK framework revealed that gritty individuals shared FC organizations among themselves in contrast to those with lower levels of grit. Such results were not observed in other behavioral measures correlated with grit such as anxiety, global impairment and social skills. With respect to network contribution, the association between FC similarity and grit was not explained by any given single functional network.

Grit was related to several behavioral measures that are important in developmental contexts. Higher grit was related to lower anxiety, general impairment and social communication skills illustrating that grit may be associated with better mental health and functioning across domains in daily life. Also of importance is that grit was not correlated with IQ, suggesting that grit isn’t simply reflecting general intellectual ability and consistent with the proposition that grit is better conceptualized as a non-cognitive trait (Duckworth et al., 2007).

Results showed that movie-related FC features (i.e., cross-movie stability, MovieTP similarity) were associated with grit, supporting the growing usage of naturalistic paradigms in the developmental research due to its suitability in individual differences fMRI research and utility in reducing head motion (Vanderwal et al., 2019; Frew et al., 2022). Though we cannot pinpoint which particular feature of MovieTP that was crucial for its association, there are several possibilities worth considering. First, we expect the context mattered more than the length of the scans as previous studies showed that longer scan duration does not necessarily yield better prediction (Vanderwal et al., 2021; Sanchez-Alonso et al., 2021). Instead, as Meer and colleagues (2020) demonstrated that subjective engagement in movie-watching sessions was related to functional dynamics, it might be possible that the emotionally engaging narrative of MovieTP ended up drawing more attention, compared to MovieDM which was an edited clip extracted from a feature-length movie. The latter might have required previous context for it to be as engaging and comprehensive as the former, which may have influenced its association with grit. We add that individual differences in attention deficit likely do not explain this interpretation due to the weak correlation found between grit and SWAN scores (Table 1). Another possibility to note is the fact that MovieTP was always presented at the tail end of the four fMRI conditions. In a developmental sample, simply staying engaged towards the end of the entire MRI session that lasted over an hour (64.7 mins) could be a burdensome task. This may have inadvertently induced a demanding situation where trait-like grit could be manifested. Although these interpretations are speculative and require more systematic examination, some combination of the movie stimuli *per se* and the grit-invoking scanning environment might have contributed to the findings.

Our analyses evinced the utility of whole-brain FCs such that no particular functional network was solely associated with grit, a finding that was corroborated by the computational lesion method in IS-RSA. Of relevance, recent studies demonstrated that FC showed greater reliability than individual edges, which implies that it may not be the simple sum of its components (Pannunzi et al., 2017; Noble et al., 2017). While it is still a possibility that specific grit-related functional networks could be revealed when using methods such as fine-scale FC (Busch et al., 2022), our findings highlight the shared whole-brain neural representations among gritty teenagers encouraging future research to examine functional systems across the brain.

Our study is not without caveats that could be addressed in future work. First, as the sample size was modest, future studies with larger samples are necessary to validate and generalize these findings. We do note that, to the extent of our knowledge, the HBN dataset is the only publicly available developmental neuroimaging dataset that includes the Grit scale as a part of its protocol. Our analyses did begin with 238 subjects from the dataset, but over 54% of the initial data had to be discarded due to excessive head motion during scanning, highlighting the challenges in procuring fMRI data with sufficient quality in developmental samples. Additionally, we focused on teenagers ranging from early adolescence to early adulthood because this is when important issues such as academic or career achievements, interpersonal relationships, mood and conduct disorders are brought to the fore (Negru-Subtirica & Pop, 2016; Kenny et al., 2013; Lee et al., 2014), coupled with dynamic neural changes encouraging the need to progress in interventions or predictions (Rosenberg et al., 2018). Future research with longitudinal study including other age groups would fill the gap by investigating whether FC features can be meaningful predictors of grit and related positive outcomes. Finally, although the present results suggest the possibility of grit-induced situations (e.g., order of scans, scan duration) being a conducive factor that captures individual differences in self-reported grit, utilizing appropriate tasks and corresponding brain-states could help circumvent the potential biases embedded within subjective behavioral assessments.

In conclusion, we provide initial evidence supporting the relationship between grit and neurodevelopmental features in adolescence and emerging adulthood. Grit was correlated with positive behavioral phenotypes such as lower anxiety, lesser general impairment, and better social skills. We showed that within-subject FC stability and between-subject FC similarity in movie-related scans were associated with grit. Notably, FC similarity while watching a complete animation film were higher amongst gritty individuals but not with others. Taken together, these results highlight potential common neural representations of grit that are distributed across the whole brain, and reveal that gritty teenagers share similar functional connectome architecture.

## Data Availability

The HBN dataset (http://fcon_1000.projects.nitrc.org/indi/cmi_healthy_brain_network/) is publicly available.

## Acknowledgements

This research was supported by the National Research Foundation of Korea (NRF-2022R1A2C1091871 and NRF-2021R1F1A1045988). We thank the original authors of the HBN dataset for their generosity in making it available for use.

## Author Contributions

S.P. and M.J.K. developed the study concept; S.P. analyzed the data under the supervision of M.J.K; S.P., D.P., and M.J.K. drafted the manuscript. All authors reviewed and approved the final manuscript for submission.

## Competing Interests

The authors declare that they have no conflict of interest.

## Notes

### Competing Interest Statement

The authors have declared no competing interest.

## References

Alexander, L. M., Escalera, J., Ai, L., Andreotti, C., Febre, K., Mangone, A., ... & Milham, M. P. (2017). An open resource for transdiagnostic research in pediatric mental health and learning disorders. Scientific Data, 4(1), 1–26.

Bird, H. R., Shaffer, D., Fisher, P., & Gould, M. S. (1993). The Columbia Impairment Scale (CIS): pilot findings on a measure of global impairment for children and adolescents. International Journal of Methods in Psychiatric Research.

Birmaher, B., Brent, D. A., Chiappetta, L., Bridge, J., Monga, S., & Baugher, M. (1999). Psychometric properties of the Screen for Child Anxiety Related Emotional Disorders (SCARED): a replication study. Journal of the American Academy of Child & Adolescent Psychiatry, 38(10), 1230–1236.

Busch, E., L., Rapuano, K. M., Anderson, K., Rosenberg, M. D., Watts, R., Casey, B. J., ... & Feilong, M. (2022). Dissociation of reliability, predictability, and heritability in fine-and coarse-scale functional connectomes during development. bioRxiv, doi:10.1101/2022.05.24.493295

Casey, B. J., & Caudle, K. (2013). The teenage brain: Self control. Current Directions in Psychological Science, 22(2), 82–87.

Casey, B. J., Somerville, L. H., Gotlib, I. H., Ayduk, O., Franklin, N. T., Askren, M. K., ... & Shoda, Y. (2011). Behavioral and neural correlates of delay of gratification 40 years later. Proceedings of the National Academy of Sciences, 108(36), 14998–15003.

Chandler, S., Charman, T., Baird, G., Simonoff, E., Loucas, T. O. M., Meldrum, D., ... & Pickles, A. (2007). Validation of the social communication questionnaire in a population cohort of children with autism spectrum disorders. Journal of the American Academy of Child & Adolescent Psychiatry, 46(10), 1324–1332.

Corriveau, A., Yoo, K., Kwon, Y. H., Chun, M. M., & Rosenberg, M. D. (2022). Functional connectome stability and optimality are markers of cognitive performance. Cerebral Cortex.

Cox, R. W. (1996). AFNI: software for analysis and visualization of functional magnetic resonance neuroimages. Computers and Biomedical Research, 29(3), 162–173.

Cox, R. W., & Hyde, J. S. (1997). Software tools for analysis and visualization of fMRI data. NMR in Biomedicine: An International Journal Devoted to the Development and Application of Magnetic Resonance In Vivo, 10(4□5), 171–178.

Dosenbach, N. U., Nardos, B., Cohen, A. L., Fair, D. A., Power, J. D., Church, J. A., ... & Schlaggar, B. L. (2010). Prediction of individual brain maturity using fMRI. Science, 329(5997), 1358–1361.

Duckworth, A., & Gross, J. J. (2014). Self-control and grit: Related but separable determinants of success. Current directions in psychological science, 23(5), 319–325.

Duckworth, A. L., Peterson, C., Matthews, M. D., & Kelly, D. R. (2007). Grit: perseverance and passion for long-term goals. Journal of Personality and Social Psychology, 92(6), 1087.

Eskreis-Winkler, L., Shulman, E. P., Beal, S. A., & Duckworth, A. L. (2014). The grit effect: Predicting retention in the military, the workplace, school and marriage. Frontiers in Psychology, 5, 36.

Esteban, O., Markiewicz, C. J., Blair, R. W., Moodie, C. A., Isik, A. I., Erramuzpe, A., ... & Gorgolewski, K. J. (2019). fMRIPrep: a robust preprocessing pipeline for functional MRI. Nature Methods, 16(1), 111–116.

Finn, E. S., Glerean, E., Khojandi, A. Y., Nielson, D., Molfese, P. J., Handwerker, D. A., & Bandettini, P. A. (2020). Idiosynchrony: From shared responses to individual differences during naturalistic neuroimaging. NeuroImage, 215, 116828.

Finn, E. S., Shen, X., Scheinost, D., Rosenberg, M. D., Huang, J., Chun, M. M., ... & Constable, R. T. (2015). Functional connectome fingerprinting: identifying individuals using patterns of brain connectivity. Nature Neuroscience, 18(11), 1664–1671.

Frew, S., Samara, A., Shearer, H., Eilbott, J., & Vanderwal, T. (2022). Getting the nod: Pediatric head motion in a transdiagnostic sample during movie-and resting-state fMRI. PLoS One, 17(4), e0265112.

Hou, X. L., Becker, N., Hu, T. Q., Koch, M., Xi, J. Z., & Mottus, R. (2022). Do grittier people have greater subjective well-being? A meta-analysis. Personality and Social Psychology Bulletin, 48(12), 1701–1716.

Horien, C., Shen, X., Scheinost, D., & Constable, R. T. (2019). The individual functional connectome is unique and stable over months to years. NeuroImage, 189, 676–687.

Jeong, J. Y., Seo, Y. S., Choi, J. H., Kim, S. H., Lee, M. S., Hong, S. H., ... & Park, D. E. (2019). The influence of grit on turnover intention of university hospital nurses: The mediating effect of job involvement. Journal of Korean Academy of Nursing, 49(2), 181–190.

Jiang, W., Xiao, Z., Liu, Y., Guo, K., Jiang, J., & Du, X. (2019). Reciprocal relations between grit and academic achievement: A longitudinal study. Learning and Individual Differences, 71, 13–22.

Kalia, V., Knauft, K. M., & Smith, A. R. (2022). Differential Associations between Strategies of Emotion Regulation and Facets of Grit in College Students and Adults. The Journal of Genetic Psychology, 183(2), 122–135.

Kaufmann, T., Alnæs, D., Doan, N. T., Brandt, C. L., Andreassen, O. A., & Westlye, L. T. (2017). Delayed stabilization and individualization in connectome development are related to psychiatric disorders. Nature Neuroscience, 20(4), 513–515.

Kelly, A. C., Di Martino, A., Uddin, L. Q., Shehzad, Z., Gee, D. G., Reiss, P. T., ... & Milham, M. P. (2009). Development of anterior cingulate functional connectivity from late childhood to early adulthood. Cerebral Cortex, 19(3), 640–657.

Kenny, R., Dooley, B., & Fitzgerald, A. (2013). Interpersonal relationships and emotional distress in adolescence. Journal of Adolescence, 36(2), 351–360.

Lam, K. K. L., & Zhou, M. (2022). Grit and academic achievement: A comparative cross-cultural meta-analysis. Journal of Educational Psychology, 114(3), 597–621.

Lee, F. S., Heimer, H., Giedd, J. N., Lein, E. S., Šestan, N., Weinberger, D. R., & Casey, B. J. (2014). Adolescent mental health—opportunity and obligation. Science, 346(6209), 547–549.

Mantel, N. (1967). The detection of disease clustering and a generalized regression approach. Cancer research, 27(2_Part_1), 209–220.

McRae, K., Gross, J. J., Weber, J., Robertson, E. R., Sokol-Hessner, P., Ray, R. D., ... & Ochsner, K. N. (2012). The development of emotion regulation: an fMRI study of cognitive reappraisal in children, adolescents and young adults. Social Cognitive and Affective Neuroscience, 7(1), 11–22.

Meer, J. N., Breakspear, M., Chang, L. J., Sonkusare, S., & Cocchi, L. (2020). Movie viewing elicits rich and reliable brain state dynamics. Nature Communications, 11(1), 1–14.

Messer, S. C., Angold, A., Costello, E. J., Loeber, R., Van Kammen, W., & Stouthamer-Loeber, M. (1995). Development of a short questionnaire for use in epidemiological studies of depression in children and adolescents: Factor composition and structure across development. International Journal of Methods in Psychiatric Research, 5, 251–262.

Moriguchi, Y., & Hiraki, K. (2013). Prefrontal cortex and executive function in young children: a review of NIRS studies. Frontiers in Human Neuroscience, 7, 867.

Myers, C. A., Wang, C., Black, J. M., Bugescu, N., & Hoeft, F. (2016). The matter of motivation: Striatal resting-state connectivity is dissociable between grit and growth mindset. Social Cognitive and Affective Neuroscience, 11(10), 1521–1527.

Negru-Subtirica, O., & Pop, E. I. (2016). Longitudinal links between career adaptability and academic achievement in adolescence. Journal of Vocational Behavior, 93, 163–170.

Nemmi, F., Nymberg, C., Helander, E., & Klingberg, T. (2016). Grit is associated with structure of nucleus accumbens and gains in cognitive training. Journal of Cognitive Neuroscience, 28(11), 1688–1699.

Noble, S., Spann, M. N., Tokoglu, F., Shen, X., Constable, R. T., & Scheinost, D. (2017). Influences on the test–retest reliability of functional connectivity MRI and its relationship with behavioral utility. Cerebral Cortex, 27(11), 5415–5429.

Pannunzi, M., Hindriks, R., Bettinardi, R. G., Wenger, E., Lisofsky, N., Martensson, J., ... & Deco, G. (2017). Resting-state fMRI correlations: From link-wise unreliability to whole brain stability. NeuroImage, 157, 250–262.

Park, D., Yu, A., Baelen, R. N., Tsukayama, E., & Duckworth, A. L. (2018). Fostering grit: Perceived school goal-structure predicts growth in grit and grades. Contemporary Educational Psychology, 55, 120–128.

Power, J. D., Fair, D. A., Schlaggar, B. L., & Petersen, S. E. (2010). The development of human functional brain networks. Neuron, 67(5), 735–748.

Robertson-Kraft, C., & Duckworth, A. L. (2014). True grit: Trait-level perseverance and passion for long-term goals predicts effectiveness and retention among novice teachers. Teachers College Record, 116(3), 1–27.

Rosenberg, M. D., Casey, B. J., & Holmes, A. J. (2018). Prediction complements explanation in understanding the developing brain. Nature Communications, 9(1), 1–13.

Salamone, J. D., & Correa, M. (2012). The mysterious motivational functions of mesolimbic dopamine. Neuron, 76(3), 470–485.

Sanchez-Alonso, S., Rosenberg, M. D., & Aslin, R. N. (2021). Functional connectivity patterns predict naturalistic viewing versus rest across development. NeuroImage, 229, 117630.

Shan, Z. Y., Mohamed, A. Z., Schwenn, P., McLoughlin, L. T., Boyes, A., Sacks, D. D., ... & Hermens, D. F. (2022). A longitudinal study of functional connectome uniqueness and its association with psychological distress in adolescence. NeuroImage, 258, 119358.

Shen, X., Tokoglu, F., Papademetris, X., & Constable, R. T. (2013). Groupwise whole-brain parcellation from resting-state fMRI data for network node identification. NeuroImage, 82, 403–415.

Swanson, J., Deutsch, C., Cantwell, D., Posner, M., Kennedy, J. L., Barr, C. L., ... & Wasdell, M. (2001). Genes and attention-deficit hyperactivity disorder. Clinical Neuroscience Research, 1(3), 207–216.

Tang, X., Wang, M. T., Guo, J., & Salmela-Aro, K. (2019). Building grit: The longitudinal pathways between mindset, commitment, grit, and academic outcomes. Journal of Youth and Adolescence, 48, 850–863.

Taylor, P. A., & Saad, Z. S. (2013). FATCAT:(an efficient) functional and tractographic connectivity analysis toolbox. Brain Connectivity, 3(5), 523–535.

Treadway, M. T., Buckholtz, J. W., Cowan, R. L., Woodward, N. D., Li, R., Ansari, M. S., ... & Zald, D. H. (2012). Dopaminergic mechanisms of individual differences in human effort-based decision-making. Journal of Neuroscience, 32(18), 6170–6176.

Valdez, J. P. M., & Datu, J. A. D. (2021). How do grit and gratitude relate to flourishing? The mediating role of emotion regulation. In Multidisciplinary Perspectives on Grit (pp. 1–16). Springer, Cham.

Vanderwal, T., Eilbott, J., Kelly, C., Frew, S. R., Woodward, T. S., Milham, M. P., & Castellanos, F. X. (2021). Stability and similarity of the pediatric connectome as developmental measures. NeuroImage, 226, 117537.

Vanderwal, T., Eilbott, J., & Castellanos, F. X. (2019). Movies in the magnet: Naturalistic paradigms in developmental functional neuroimaging. Developmental Cognitive Neuroscience, 36, 100600.

Wang, S., Dai, J., Li, J., Wang, X., Chen, T., Yang, X., ... & Gong, Q. (2018). Neuroanatomical correlates of grit: Growth mindset mediates the association between gray matter structure and trait grit in late adolescence. Human Brain Mapping, 39(4), 1688–1699.

Wang, S., Zhou, M., Chen, T., Yang, X., Chen, G., Wang, M., & Gong, Q. (2017). Grit and the brain: spontaneous activity of the dorsomedial prefrontal cortex mediates the relationship between the trait grit and academic performance. Social Cognitive and Affective Neuroscience, 12(3), 452–460.

Wechsler, D. (2014). WISC-V: Technical and interpretive manual. NCS Pearson, Incorporated.

Wolters, C. A., & Hussain, M. (2015). Investigating grit and its relations with college students’ self-regulated learning and academic achievement. Metacognition and Learning, 10, 293–311.

