## Supplemental Materials for "Similarity in Functional Connectome Architecture Predicts Teenage Grit"

### Supplemental Methods

#### *Participants*

Sample characteristics of the final sample of 64 participants analyzed in this study are presented in Figure S1 (22 females; age range = 11.07-18.82 years ($M$ = 14.78); $n$ = 48 from CBIC).

**A.** **B.**

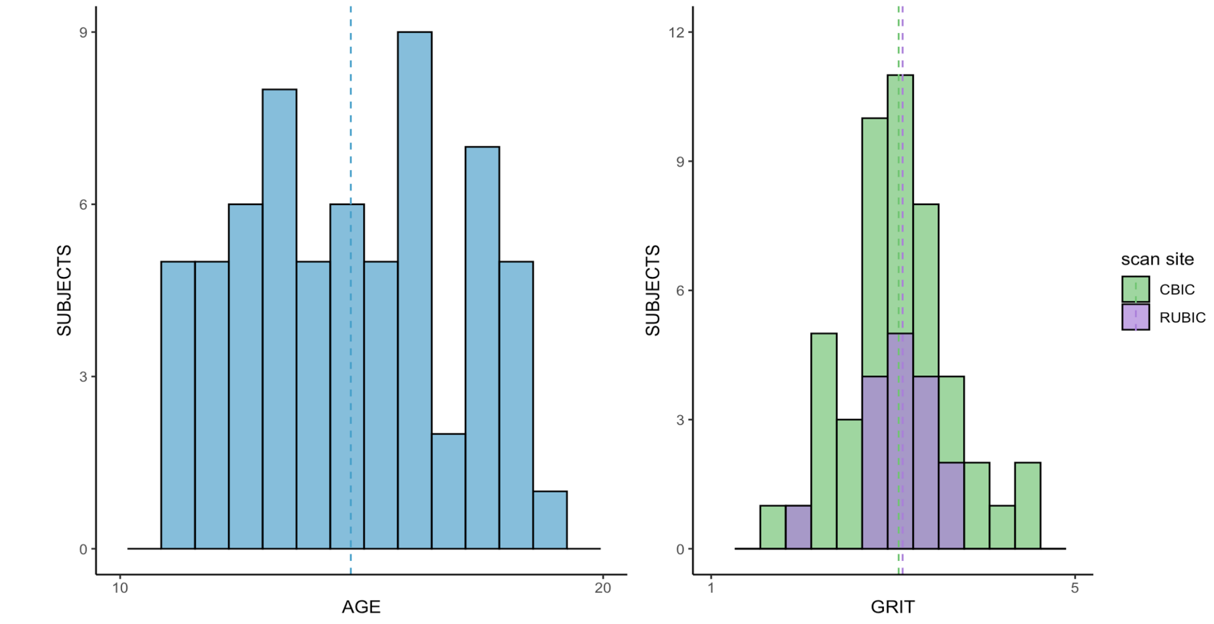


**Figure S1**. Sample characteristics ($N$ = 64). **A.** Age distribution of the sample used for this study. Mean age (14.78 years) is presented in a dashed line. **B.** Distributions of grit scores for the two scan sites are presented. Dashed lines are mean grit score, 3.06 for CBIC and 3.10 for RUBIC.

### Supplemental Results

*Correlations*

The correlations between stability and similarity measures and chronological age of the sample are presented in Table S1. None of the six measures were significantly correlated with age. The six functional connectome measures within each of the 10 canonical networks in MovieTP condition were not significantly correlated with total grit score (Table S2).

**Table S1.** Results of Correlations between FC Measures and Participants' Chronological Age.

| | | within-subject FC similarity | | | between-subject FC similarity | | |
|---|---|---|---|---|---|---|---|
| | | cross-rest | cross-state | cross-movie | Rest1 | MovieDM | MovieTP |
| Full Volume | | | | | | | |
| | age | .21 (.0960) | .03 (.8388) | - .01 (.9490) | .03 (.7981) | .02 (.8778) | .02 (.8888) |
| Volume-matched | | | | | | | |
| | age | .15 (.2300) | - .02 (.8577) | - .14 (.2687) | .01 (.9296) | - .10 (.4282) | .02 (.8888) |

Note. $N$ = 64, Partial correlation coefficients ($r$) and corresponding $p$-values in parenthesis (Bonferroni corrected, $p <$ .008).

**Table S2.** Results of Correlations between FC Measures of each 10 Canonical Networks in MovieTP Condition and Total Grit Score.

| Medial Frontal | Fronto parietal | Default mode | Motor | Visual I | Visual II | Visual Association | Limbic | Basal Ganglia | Cerebellum |
|---|---|---|---|---|---|---|---|---|---|
| .03 (.7911) | .17 (.1940) | .24 (.0699) | .18 (.1794) | .10 (.4679) | .07 (.6194) | - .02 (.8542) | .17 (.1994) | - .04 (.7565) | .02 (.9059) |

Note. $N$ = 64, Partial correlation coefficients ($r$) and corresponding $p$-values in parenthesis (Bonferroni corrected, $p <$ .005).

*Functional connectivity map*

Group-averaged functional connectivity patterns of high and low grit groups (median-split) are summarized in Figure S2.

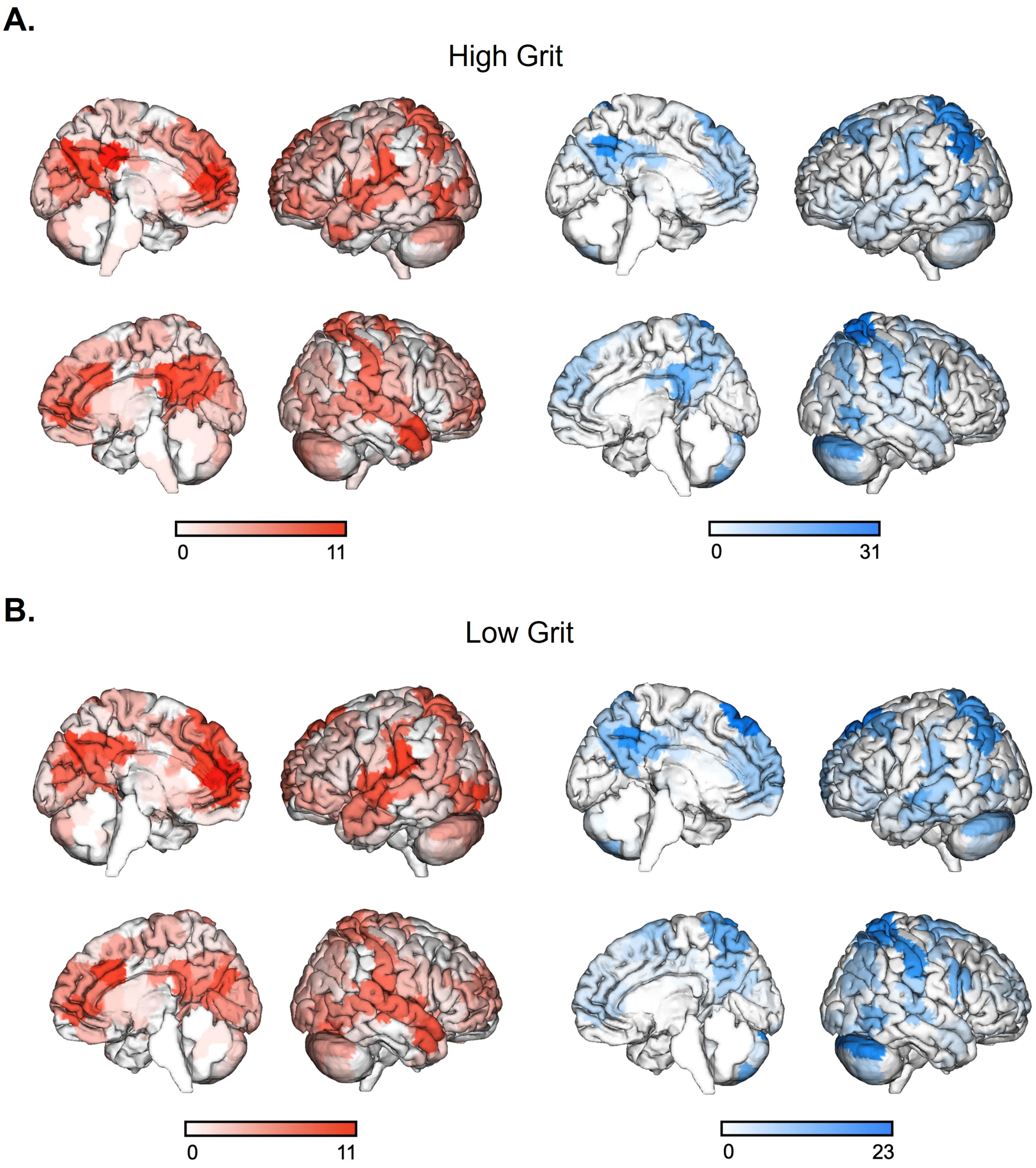


**Figure S2**. Visualization of positive (red) and negative (blue) functional connectivity patterns during movie-watching in individuals with high and low grit. For visualization purposes, edges with top 1% highest positive and negative functional connectivity values are shown. Color bars denote the maximum number of functional connectivity for each node.
